# Environmental NaCl affects *C. elegans* development and aging

**DOI:** 10.1101/2025.03.09.641258

**Authors:** Franziska Pohl, Brian M. Egan, Daniel L. Schneider, Matthew C. Mosley, Micklaus A. Garcia, Sydney Hou, Chen-Hao Chiu, Kerry Kornfeld

## Abstract

Sodium is an essential nutrient, but is toxic in excess. In humans, excessive dietary sodium can cause high blood pressure, which contributes to age-related diseases including stroke and heart disease. We used *C. elegans* to elucidate how sodium levels influence animal aging. Most experiments on this animal are conducted in standard culture conditions: Nematode Growth Medium (NGM) agar with a lawn of *E. coli*. Here, we report that the supplemental NaCl in standard NGM, 50 mM, accelerates aging and decreases lifespan. For comparison, we prepared NGM with reduced NaCl or excess NaCl. Considering reduced NaCl as a baseline, wild-type worms on standard NGM displayed normal development and fertility but reduced lifespan and health span, indicating toxicity in old animals. The long-lived mutants *daf-*2, *age-1*, and *nuo-6,* cultured on standard NGM, also displayed reduced lifespan. Thus, NaCl in standard NGM accelerates aging in multiple genetic backgrounds. Wild-type worms on excess NaCl displayed delayed development and reduced fertility, and reduced lifespan and health span, indicating toxicity in both young and old animals. These results suggest that young animals are relatively resistant to NaCl toxicity, but that aging causes progressive sensitivity, such that old animals display toxicity to both standard and excess NaCl. We investigated pathways that respond to NaCl. Young animals cultured with excess NaCl activated *gpdh-1,* a specific response to NaCl stress. Old animals cultured with excess NaCl activated *gpdh-1* and *hsp-6*, a reporter for the mitochondrial unfolded protein response. Thus, excess NaCl activates multiple stress response pathways in older animals.

## Introduction

Standard conditions used to culture *C. elegans* in the laboratory are quite different from their natural environment (Félix and Braendle 2010; Guisnet et al. 2021). Wild worms live on rotting plant material and may experience rapid and extreme changes in their physical, chemical, and microbiological environment (Félix and Duveau 2012; Schulenburg and Félix 2017). Laboratory *C. elegans* are typically cultured on nematode growth medium (NGM), maintained at a constant temperature of 20°C, and provided excess food in the form of *E. coli* OP50 bacteria. Laboratory worms rarely experience extreme stress and spend most of their lifespan as adults. These “standard culture conditions” were established by Sydney Brenner to study genetic mutations that affect the nervous system or development (Brenner 1974; Sulston et al. 1983). They have been widely adopted, which facilitates comparisons between laboratories. However, standard culture conditions were not optimized for the long-term survival of individuals; unsurprisingly, several aspects of standard culture conditions have since been demonstrated to influence aging and lifespan, including temperature, pathogenicity of *E. coli*, and oxygen levels (Klass 1977; Adachi et al. 1998; Garigan et al. 2002; Huang et al. 2004; Yu et al. 2015). Because *C. elegans* has become a leading model for studies of aging and lifespan, it is important to understand whether additional features of standard culture conditions influence aging.

Sodium is an essential nutrient, but it can be toxic in excess. In humans, excessive ingestion of dietary sodium is a prevalent cause of high blood pressure, which contributes to strokes and heart disease that shorten lifespan. Here, we focus on how supplemental NaCl added to NGM agar affects aging. Animals have evolved mechanisms to sense and regulate osmolarity in response to variable environments or diets (Bourque 2008). *C. elegans* osmoregulate by sensing and responding to high osmolarity (Choe 2013) via osmoregulatory organs, including the hypodermis, intestine, excretory cell, and amphid neurons (Huang et al. 2007). To characterize interactions between aging and osmoregulation in worms, we varied the level of supplemental NaCl in NGM. Standard NGM contains 50 mM supplemental NaCl in addition to undefined levels of NaCl contributed by other components. We prepared NGM without supplemental NaCl (reduced NaCl medium) and with 200 mM supplemental NaCl (excess NaCl medium). Excess NaCl was toxic for young and old wild-type animals, causing delayed development, reduced fecundity, and reduced lifespan and health span. Standard NaCl was benign in young animals but toxic in old animals, reducing lifespan and health span, indicating that the NaCl added to standard NGM accelerates aging. We replicated the lifespan reduction caused by standard NaCl in three long-lived mutant strains. To investigate how animals respond to NaCl, we measured stress response pathways. Young animals cultured with excess NaCl displayed transcriptional activation of *gpdh-1,* a specific response to NaCl stress. Old animals cultured with excess NaCl displayed transcriptional activation of *gpdh-1* and *hsp-6*, a reporter for the mitochondrial unfolded protein response. Thus, excess NaCl activates multiple stress response pathways in older animals. These results suggest that young animals are relatively resistant to NaCl toxicity, thriving in reduced and standard NaCl levels and only displaying toxicity in excess NaCl. Aging causes progressive sensitivity to NaCl toxicity, so that old animals display toxicity in standard and excess NaCl. Furthermore, nearly all previous studies of *C. elegans* using standard NGM were performed under conditions of mild osmotic stress that are tolerable to young animals but detrimental to old animals.

## Results

### Young animals tolerate standard NaCl but display toxicity in excess NaCl, whereas old animals display toxicity in both standard and excess NaCl

To investigate the effects of environmental NaCl on aging, we quantified the lifespan of wild-type worms cultured from the egg stage on standard NGM (50 mM supplemental NaCl), NGM with excess NaCl (200 mM supplemental NaCl), and NGM with reduced NaCl (0 mM supplemental NaCl). Excess NaCl significantly decreased mean lifespan to ∼12 days, compared to ∼16 days in standard NGM (**Fig.1A**), consistent with previous reports (Lamitina et al. 2004; Chandler-Brown et al. 2015). However, reduced NaCl significantly increased mean lifespan to ∼18 days, suggesting that this concentration of NaCl extends lifespan – or, conversely, that the concentration of NaCl in standard NGM shortens lifespan (**Fig.1A**). To investigate health span, we quantified the age-related declines in pharyngeal pumping and body movement. Compared to standard NGM, excess NaCl accelerated the age-related decline of pharyngeal pumping (**Fig.1B**). Reduced NaCl significantly increased pharyngeal pumping at adult days 6-8 compared to standard NGM, suggesting that the concentration of NaCl in standard NGM decreases pharyngeal pumping health span. We evaluated body movement by two methods: counting body bends per minute, and categorizing animals as class A (robust movement), class B (moderate impairment), or class C (severe impairment). Compared to standard NGM, excess NaCl significantly reduced the number of body bends per minute at days 4-9, whereas reduced NaCl increased body bends per minute at day 8 (**Fig.1C**). Based on movement class, excess NaCl accelerated age-related decline in movement, whereas reduced NaCl delayed age-related decline: day 8 animals cultured on reduced NaCl were 72% class A, compared to 47% for standard NGM, and 20% for excess NaCl (**Fig.1D**). A similar trend was observed from days 6-11 (**Fig.S1A-C**). These results suggest that the concentration of NaCl in standard NGM decreases body movement health span. *C. elegans* display an age-related increase in autofluorescence (Pincus et al. 2016; Komura et al. 2021). Excess and reduced NaCl did not significantly affect the level of autofluorescence intensity on adult days 0, 5, and 11 (**Fig.S1E-F**).

**Figure 1.**
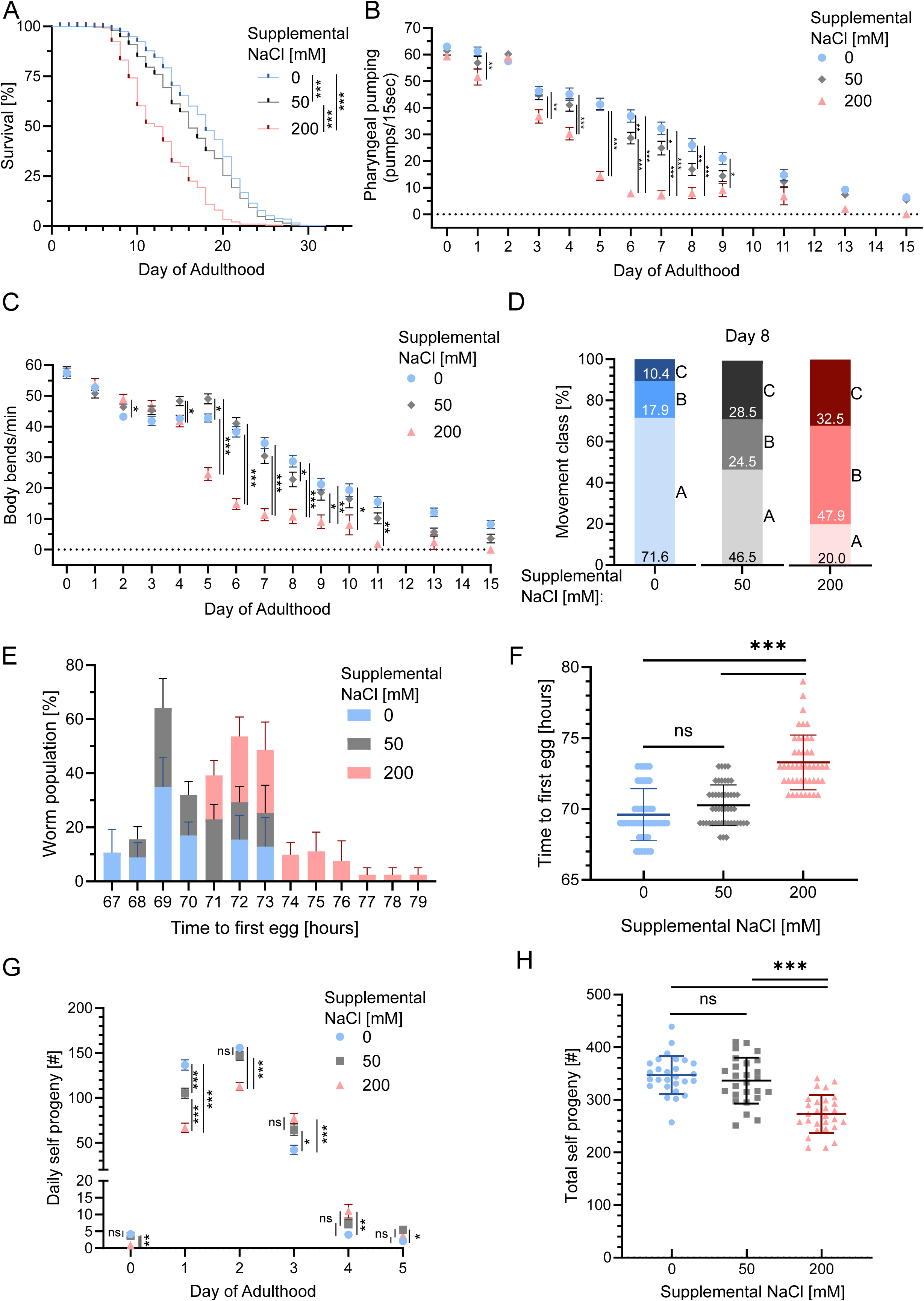
Environmental NaCl affects wild-type *C. elegans* lifespan, health span, development rate, and progeny production. **A)** Hermaphrodite embryos were cultured on NGM dishes with the indicated concentration of supplemental NaCl. The resulting adults were transferred to fresh dishes periodically to separate them from progeny and monitored daily for survival beginning on adult day 0. Eight independent experiments were performed with a combined 625, 584, 223 death events, and 122, 154, 159 censored subjects for 0, 50, 200 mM supplemental NaCl, respectively. Kaplan-Meier Analysis with log-rank test for trend. p-values were corrected using Bonferroni correction. ***p<0.001. **B,C)** Pharyngeal pumping and body bends were observed with a dissecting microscope. Three independent experiments with ≥15 animals per experiment. Values are mean±SEM analyzed using ordinary two-way ANOVA with Tukey’s multiple comparison test, with a single pooled variance. *p<0.05, **p<0.01, ***p<0.001. **D)** Movement classes on day 8 were categorized with a dissecting microscope; bars with shaded colors display percent of the population in class A on bottom, B in the middle, and C on top. Two independent experiments with a combined n=76, 62, and 27 animals cultured on 0, 50, and 200 mM supplemental NaCl, respectively. Class A: vigorous, coordinated, spontaneous movement; Class B: uncoordinated movement - part of body (head, tail, etc.) is paralyzed or uncoordinated; Class C: majority of animal is paralyzed, no forward or backward movement, only moves head or tail slightly. **E,F)** Animals were synchronized at the egg stage and monitored hourly for deposition of their first egg, resulting in one value per animal. Three independent experiments with ≥7 animals per experiment. Bars in **E** are mean±SEM; bars in **F** are average time to first egg, mean±SD. Ordinary one-way ANOVA with Dunnett’s multiple comparison test with single pooled variance. ***p<0.001. **G,H)** Day 0 adult hermaphrodites were transferred to new dishes daily, and their daily progeny production was measured by counting their progeny 2-5 days later. Three independent experiments with ≥5 hermaphrodites. **G)** Daily progeny production shown as mean±SEM, analyzed using mixed-effects model with Geisser-Greenhouse correction and Tukey’s multiple comparison test, with individual variances computed for each comparison. **F)** Total number of self-progeny for day 0-5 of individual animals, shown as mean±SD, analyzed using ordinary one-way ANOVA and Dunnett’s multiple comparison test with a single pooled variance. *p<0.05, **p<0.01, ***p<0.001.

To evaluate the effect of NaCl on developmental rate, we quantified the time between an egg hatching and the sexually mature hermaphrodite laying its first egg. On reduced NaCl, hermaphrodites generated their first egg after 67-73 hours, similar to animals on standard NGM (68-73 hours). By contrast, on excess NaCl, hermaphrodites displayed a significantly delayed time to first egg of 71-79 hours (**Fig.1E-F**). To evaluate progeny production, we measured daily and total self-progeny of wild-type hermaphrodites. Compared to standard NGM, reduced NaCl slightly increased progeny production on adult day 1 and slightly reduced progeny production on adult days 3 and 5 (**Fig**.**1G**); the total number of self-progeny was not affected (**Fig.1H**). By contrast, excess NaCl significantly reduced progeny production on days 0-2 (**Fig**.**1G**), and the total number of self-progeny was significantly reduced (**Fig**.**1H**). Thus, for young animals, standard NaCl has a minimal effect on developmental rate and fertility, whereas excess NaCl delays developmental rate and reduces overall fecundity.

Standard NGM contains 50 mM supplemental NaCl, but it also contains other ingredients that contain NaCl in unknown quantities. Thus, when we omit 50 mM supplemental NaCl from NGM, NaCl in the medium is reduced but not eliminated. To evaluate the contribution of NaCl from other ingredients, we used inductively coupled plasma mass spectrometry (ICP-MS) to analyze the elemental composition of agar, peptone, yeast extract, and tryptone. Peptone and agar are ingredients of NGM (Brenner 1974) whereas yeast extract and tryptone are ingredients of LB that is used to initially culture *E. coli* for consumption by *C. elegans.* Tryptone displayed the highest concentration of sodium (27,900 ± 805 parts per million (ppm)), followed by peptone (17,200 ± 360 ppm), agar (10,800 ± 500 ppm), and yeast extract (4,840 ± 590 ppm) (**Fig.S2B**). ICP-MS was also used to determine the concentrations of other elements, and **Figure S2C-E** presents values for potassium, magnesium, calcium, iron, zinc, vanadium, chromium, cobalt, and other elements. To determine if evaporative loss affects the concentration of NaCl in NGM dishes, we measured the loss of mass over time at 4°C and at 20°C (**Fig.S2F,G**). At 4°C, the mean mass of NGM dishes with 0, 50, or 200 mM supplemental NaCl remained at >95% of their original mass over a period of 7 weeks (49 days). By contrast, at 20°C, the mass of NGM dishes was reduced more quickly, dropping below 95% of their original mass at day 17. NaCl concentration did not significantly affect evaporation at 4°C, but did at 20°C. To minimize the effect of evaporation during our experiments, we stored NGM dishes at 4°C for no longer than 2 months and at 20°C for no longer than 7 days.

### Lifespan extensions caused by reduced NaCl and insulin/IGF signaling mutations are additive

To investigate the lifespan extension caused by reduced NaCl, we analyzed mutations that alter longevity. Mutation of insulin/insulin-like growth factor (IGF) signaling pathway members controls lifespan, either extending it (*daf-2, age-1,* etc.) or reducing it (*daf-16, daf-12,* etc.) (Friedman and Johnson 1988; Kenyon et al. 1993; Kimura et al. 1997; Ogg and Ruvkun 1998; Paradis and Ruvkun 1998). *daf-2(e1370)* and *age-1(hx546)* mutants cultured on reduced NaCl displayed significantly increased mean lifespans (**Fig.2A-B**, **Table 1**). Thus, the lifespan extension caused by reduced NaCl was additive with the lifespan extensions caused by reduced insulin/IGF signaling, indicating that reduced NaCl likely extends lifespan via a distinct mechanism. Culturing these mutants in excess NaCl medium decreased lifespan in both mutant strains. Thus, these mutations do not confer resistance to high NaCl stress. By contrast, *daf-12* and *daf-16* mutants cultured on reduced NaCl did not display significantly increased lifespan compared to standard NGM, whereas culture in excess NaCl significantly decreased lifespan (**Fig.2C-D**, **Table 1**).

**Figure 2.**
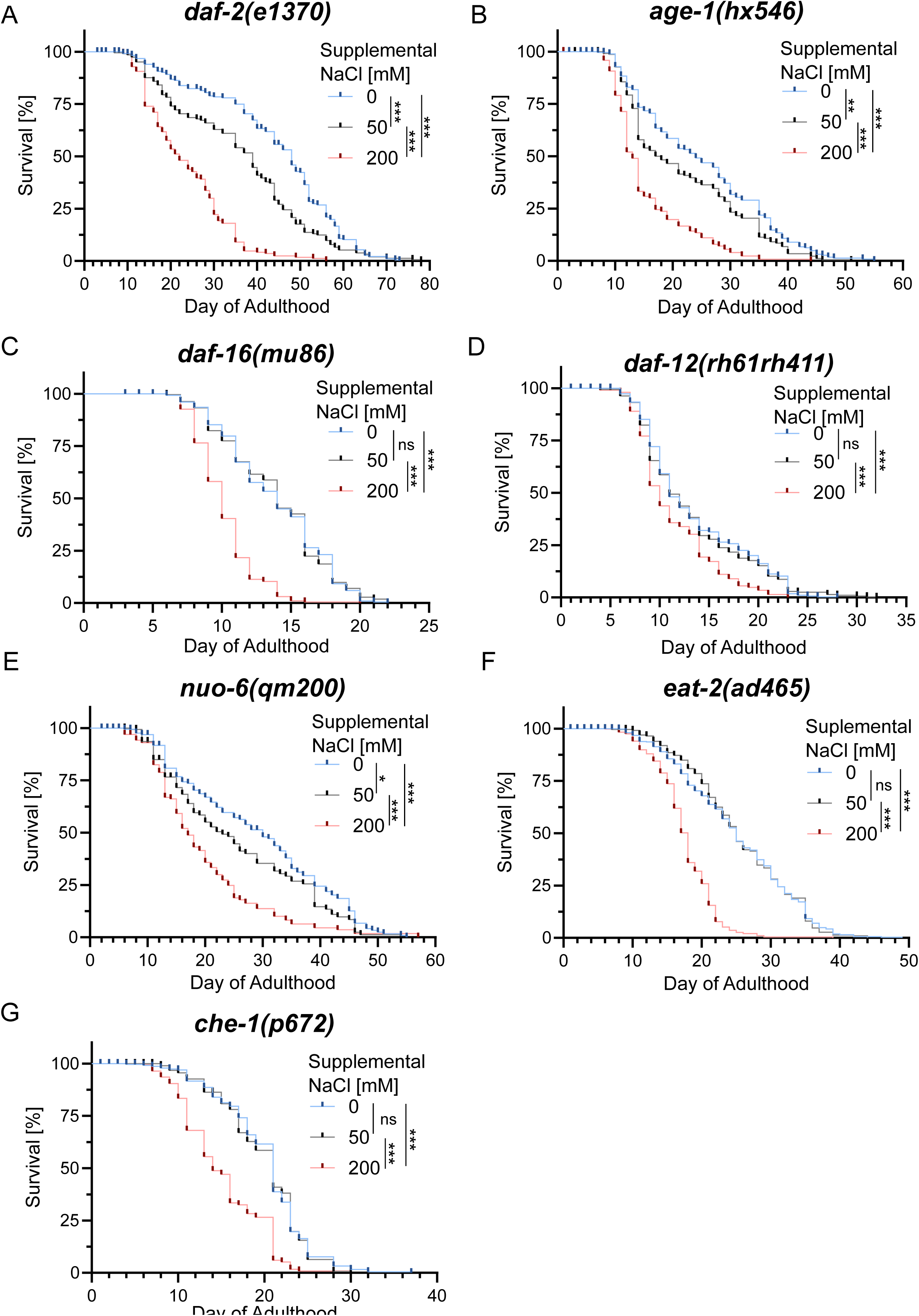
Environmental NaCl influences the lifespan of mutant *C. elegans*. Hermaphrodite embryos were cultured on NGM dishes with the indicated concentration of supplemental NaCl, and the resulting adults were monitored daily for survival beginning on adult day 0. Survival curves were analyzed by Kaplan-Meier Analysis with log-rank test for trend; p-values were corrected using Bonferroni correction. *p<0.05 **p<0.01, ***p<0.001. **A)** *daf-2(e1370)*: Four independent experiments with 195, 162, 172 death events and 55, 60, 87 censored subjects for 0, 50, 200 mM supplemental NaCl, respectively. **B)** *age-1(hx546)*: Three independent experiments with 235, 182, 146 death events and 56, 103, 150 censored subjects for 0, 50, 200 mM supplemental NaCl, respectively. **C)** *nuo-6(qm200)*: Three independent experiments with 123, 93, 130 death events and 103, 132, 93 censored subjects for 0, 50, 200 mM supplemental NaCl, respectively. **D)** *eat-2(ad465)*: Four independent experiments with 367, 322, 236 death events and 134, 173, 154 censored subjects for 0, 50, 200 mM supplemental NaCl, respectively. **E)** *daf-16(mu86)*: Three independent experiments with 237, 219, 215 death events and 27, 41, 44 censored subjects for 0, 50, 200 mM supplemental NaCl, respectively. **F)** *daf-12(rh61rh411)*: Four independent experiments with 296, 286, 186 death events and 56, 54, 141 censored subjects for 0, 50, 200 mM supplemental NaCl, respectively. **G)** *che-1(p672)*: three independent experiments with 187, 146, 135 death events and 66, 97, 107 censored subjects for 0, 50, 200 mM supplemental NaCl, respectively.

**Table 1.** Median and mean lifespan and percent change in mean survival in worms cultured on reduced NaCl (0 mM), standard NGM (50 mM), and excess NaCl (200 mM) medium.

| Genotype | Supplemental NaCl [mM] | Median survival | Mean survival (days±S.D.) | % Change mean survival | Significance vs 50 mM NaCl | n-death events |
| --- | --- | --- | --- | --- | --- | --- |
| Wild type (WT) | 0 | 18 | 17.76±5.079 | +8.1 | *** | 625 |
|  | 50 | 16 | 16.43±5.227 |  |  | 584 |
|  | 200 | 12 | 12.18±4.297 | -25.9 | *** | 223 |
| *daf-2(e1370)* | 0 | 48 | 43.03±15.75 | + 21.9 | *** | 195 |
|  | 50 | 39 | 35.31±15.88 |  |  | 162 |
|  | 200 | 22 | 23.36±9.812 | -33.8 | *** | 172 |
| *age-1(hx546)* | 0 | 24 | 24.46±11.44 | +13.9 | ** | 235 |
|  | 50 | 17 | 21.48±10.54 |  |  | 182 |
|  | 200 | 13 | 14.75±6.65 | -31.3 | *** | 146 |
| *daf-16(mu86)* | 0 | 14 | 13.77±3.703 | ±0 | ns | 237 |
|  | 50 | 14 | 13.77±3.792 |  |  | 219 |
|  | 200 | 10 | 10.05±2.112 | -27.0 | *** | 215 |
| *daf-12(rh61rh411)* | 0 | 11 | 13.17±5.271 | +1.5 | ns | 296 |
|  | 50 | 11 | 12.98±5.471 |  |  | 286 |
|  | 200 | 10 | 10.69±3.765 | -17.6 | *** | 186 |
| *nuo-6(qm200)* | 0 | 30 | 28.37±13.11 | +18.2 | * | 123 |
|  | 50 | 23 | 24.01±12.42 |  |  | 93 |
|  | 200 | 17 | 18.69±9.722 | -22.2 | *** | 130 |
| *eat-2(ad465)* | 0 | 25 | 24.34±8.392 | -1.9 | ns | 367 |
|  | 50 | 25 | 24.80±7.728 |  |  | 322 |
|  | 200 | 18 | 16.94±4.435 | -31.7 | *** | 236 |
| *che-1(p672)* | 0 | 21 | 19.94±5.270 | +1.5 | ns | 187 |
|  | 50 | 21 | 19.65±4.979 |  |  | 146 |
|  | 200 | 14 | 14.41±4.851 | -26.7 | *** | 135 |

Mutations that reduce mitochondrial function can extend lifespan (Ewbank et al. 1997; Felkai et al. 1999; Feng et al. 2001; Dillin et al. 2002). *nuo-6* encodes a conserved subunit of mitochondrial complex I (NUDFB4)(Yang and Hekimi 2010a), and a loss-of-function mutation significantly extends lifespan. *nuo-6(qm200)* mutants cultured on reduced NaCl displayed significantly extended lifespan (**Fig.2E**). *nuo-6(qm200)* mutants cultured on excess NaCl had a significantly reduced lifespan. Thus, the lifespan extension caused by reduced NaCl was additive with the lifespan extension caused by reduced mitochondrial function.

Dietary restriction is well-established to extend lifespan in many animals, including *C. elegans*. Mutations in *eat-2* impair pharyngeal pumping, allowing live *E. coli* bacteria into intestine, activating an innate immune response that includes bacterial avoidance behavior, and inducing dietary restriction (Lakowski and Hekimi 1998; McKay et al. 2004; Kumar et al. 2019). *eat-2(ad465)* mutants cultured on reduced NaCl did not display a significantly altered lifespan compared to standard NGM; mutants cultured on excess NaCl had a significantly reduced lifespan (**Fig.2F**, **Table 1**). Thus, the lifespan extensions caused by reduced NaCl and dietary restriction were not additive.

To investigate the connection between aging and mechanisms that sense osmolarity, we analyzed a *che-1* mutant. CHE-1 is a transcription factor that induces the expression of salt sensing ASE- specific target genes (Kunitomo et al. 2013; Traets et al. 2021). *che-1(p672)* mutants cultured on reduced NaCl did not display significantly altered lifespan compared to standard NGM; mutants cultured on excess NaCl had a significantly reduced lifespan (**Fig.2G**, **Table 1**). Thus, the lifespan extensions caused by reduced NaCl medium may involve salt sensing by the ASE neuron.

### Excess NaCl stimulates specific stress response pathways

Our results suggest that standard NaCl is tolerated in young animals but is toxic in old adults, whereas excess NaCl is toxic in young and old animals. To investigate pathways that might respond to NaCl stress, we analyzed reporter strains throughout adult life. We first analyzed a stress response pathway that responds specifically to excess NaCl. Worms confronted with high osmolarity increase synthesis of the organic osmolyte glycerol, a small polyol that accumulates to high levels and protects against osmotic stress (Lamitina et al. 2004; Lamitina et al. 2006; Urso et al. 2020; Urso and Lamitina 2021). *gpdh-1 and gpdh-2* (glycerol-3-phosphate dehydrogenase-1/2) encode enzymes that catalyze the rate-limiting step of glycerol biosynthesis. *gpdh-1* exhibits a strong and sustained transcriptional up-regulation during hypertonic stress, whereas *gpdh-2* is weakly and transiently up-regulated. Induction of *gpdh-1* occurs rapidly, within the first hour of exposure, and is specific to osmotic stress, since it does not occur during heat, oxidative, or endoplasmic reticulum stress. To evaluate this response, we monitored *gpdh-1* expression levels throughout the adult lifespan using two *gdph-1* GFP reporters, which gave similar results (Lamitina et al. 2006; Urso et al. 2020). Animals cultured on reduced and standard NaCl medium displayed a similar pattern; GFP expression gradually increased from day 0 to 16 of adulthood, indicating weak pathway activation in older animals (**Fig.3**). Excess NaCl caused a significant increase in expression in young adults (day 0 and 2). On adult days 4-8, expression declined and was not significantly different compared to standard NGM. Expression began to increase again on adult day 10 and was significantly and progressively increased until day 16 (**Fig.3**). Thus, young adults confronted with excess NaCl strongly induce expression of *gpdh-1*, which is likely to protect against osmotic stress. The response diminishes in middle aged adults, perhaps because homeostasis has been reestablished. The response is reactivated in older adults, perhaps because aging makes animals more vulnerable to excess NaCl stress.

**Figure 3.**
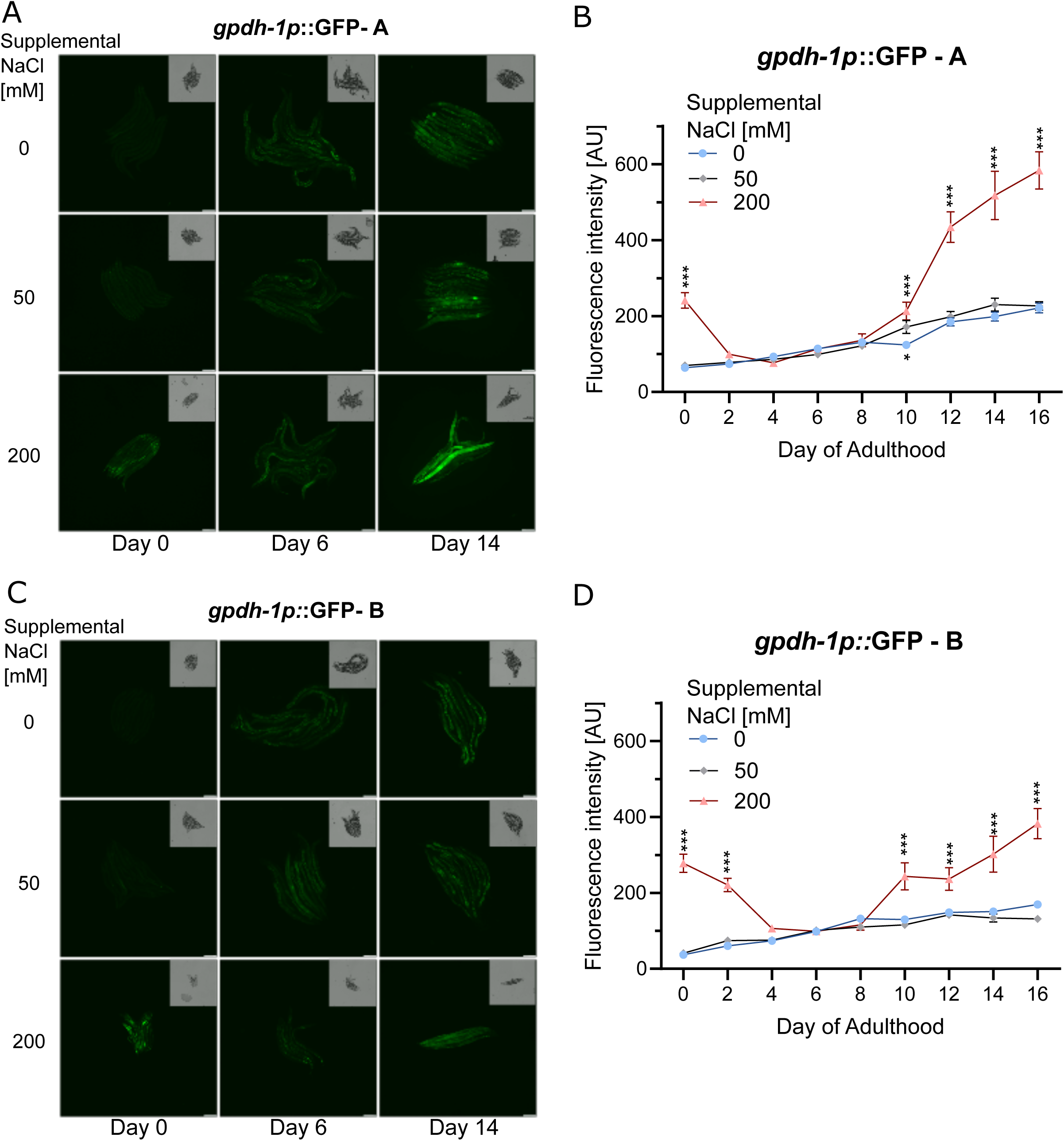
Environmental NaCl influences *gpdh-1*p expression. **A-D)** Transgenic animals containing reporter constructs were cultured on 0, 50, and 200 mM supplemental NaCl, and GFP fluorescence was quantified on adult days 0, 2, 4, 6, 8, 10, 12, 14, and 16. **A,C)** Representative fluorescence images of ∼10 animals on adult day 0, 6, and 14 cultured on medium with 0, 50, or 200 mM supplemental NaCl. Genotypes are *gpdh-1p*::GFP – A (OG119; *drIs4* [*gpdh-1p*::GFP + *col-12p*::DsRed] IV) in panels A-B, or *gpdh-1p*::GFP – B (VP198; kbIs5 [*gpdh-1p*::GFP + rol-6(su1006)]) in panels C-D. Inlay is bright field image. Scale bar on bottom right = 200 µm**. B,D)** Data points represent fluorescence intensity in arbitrary units (AU); bars are mean±SEM. Analyzed by two-way ANOVA with Dunnett’s multiple comparison analysis, with a single pooled variance. *p<0.05, ***p<0.001. For panel B, three independent experiments with n= 3-14 animals per condition each day; for panel D, three independent experiments for days 0-14, and two independent experiments for day 16, with n= 3-15 animals per condition each day.

We next examined more general stress response pathways. The mitochondrial unfolded protein response (mitoUPR)(Moehle et al. 2019; Bar-Ziv et al. 2020; Bar-Ziv et al. 2020) is induced by misfolded proteins in the mitochondria (Berendzen et al. 2016), mutations in mitochondrial genes (Lin et al. 2016; Dues et al. 2017; Wu et al. 2018), reactive oxygen species (Runkel et al. 2013; Shao et al. 2016; Pohl et al. 2023), ethidium bromide (Yoneda et al. 2004), and heat (Yu et al. 2024). The response is characterized by activation of mitochondrial chaperons such as *hsp-6* and *hsp-60*, which can be measured using an *hsp-6p*::GFP reporter strain*. hsp-6p*::GFP was expressed at a low level on adult days 0 and 5 that was not affected by reduced or excess NaCl; expression levels increased on adult day 11, and this increase was significantly enhanced by excess NaCl (**Fig.4A**). Thus, excess NaCl may damage mitochondria in old adults, stimulating the mitoUPR.

**Figure 4.**
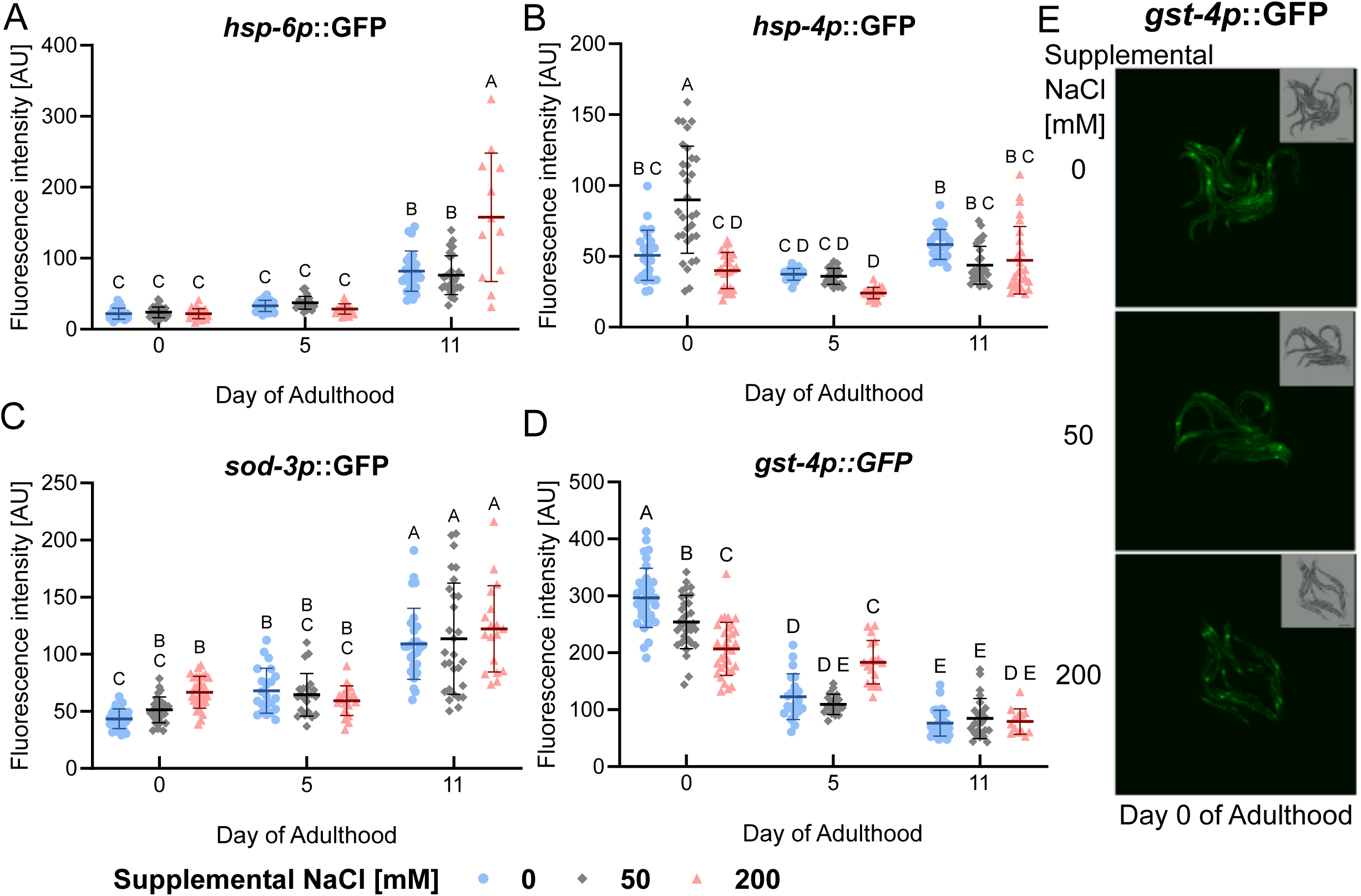
Environmental NaCl influences stress response pathways. **A-D)** Transgenic animals containing reporter constructs were cultured on 0, 50, and 200 mM supplemental NaCl, and GFP fluorescence was quantified on days 0, 5, and 11 of adulthood; each data point represents fluorescence intensity of one animal in arbitrary units (AU); bars are mean±SD. Two-way ANOVA with Tukey’s multiple comparisons test was performed using the compact letter display for comparison: for any two groups that share a letter, p>0.05, indicating no significant difference; for groups with different letters, p<0.05, indicating a statistically significant difference. **A)** Strain SJ4100 (*zcIs13 [hsp-6p::GFP + lin-15(+)]*) uses expression of GFP by the *hsp-6* promoter to report on the mitochondrial unfolded protein response (mitoUPR). **B)** Strain SJ4005 *(zcIs4 [hsp-4::GFP]* V) uses expression of *hsp-4* to report on the endoplasmic reticulum (ER) unfolded protein response. **C)** Strain CF1553 (*muIs84 [(pAD76) sod-3p::GFP + rol-6(su1006)])* uses expression of manganese superoxide dismutate 3 (*sod-3*) to report on the oxidative stress response. **D)** Strain CL2166 (*dvIs19 [(pAF15)gst-4p::GFP::NLS*] uses expression of glutathione S-transferase 4 (*gst-4*) to report on the oxidative stress response (conjugation of reduced glutathione to exogenous and endogenous hydrophobic electrophiles). **E)** Representative fluorescence images of *gst-4p::GFP* expression on day 0 from ∼10 animals cultured on NGM with 0, 50, and 200 mM supplemental NaCl. Inlay is bright field image. Scale bar on bottom right = 200 µm.

The ER unfolded protein response can be activated by heat, caffeine (Al-Amin et al. 2016), and tunicamycin (Mörck et al. 2009); it can be monitored with an *hsp*-*4p*::GFP reporter strain. Day 0 adults cultured on reduced or excess NaCl displayed reduced expression of *hsp*-*4p*::GFP compared to those cultured on standard NGM; there were no significant effects of excess or reduced NaCl on adult days 5 and 11 (**Fig.4B**). Superoxide dismutase is an enzyme that converts superoxide to hydrogen peroxide and oxygen to reduce oxidative stress (Sheng et al. 2014). A *sod*-*3p*::GFP reporter strain is activated by environmental stressors such as fungi (Kitisin et al. 2022), glucose (Kingsley et al. 2021), oxidative stress induced by paraquat, and mutations in mitochondrial genes (Schaar et al. 2015; Dues et al. 2017; Li et al. 2024). Here, *sod-3p*::GFP displayed a significant increase of expression over time, but reduced and excess NaCl had minimal effects (**Fig.3C**).

Drug-metabolizing enzymes called glutathione S-transferases utilize glutathione in conjugation reactions involved in phase II detoxification. *gst-4* is activated by multiple environmental stressors such as xenobiotics (Ferguson and Bridge 2019) and juglone (Stefanello et al. 2015). To monitor this response, we used a *gst-4p::GFP* reporter strain. Expression levels were high in day 0 animals and generally decreased on adult days 5 and 11 (**Fig.4D**). *gst-4* mRNA expression levels determined by RNA-seq also displayed this general trend (**Fig.S1G**). In day 0 adults, reduced NaCl caused significantly increased GFP expression, whereas excess NaCl caused significantly decreased GFP expression (**Fig.4D-E**).

## Discussion

When Sydney Brenner first developed *C. elegans* as model organism, he chose culture conditions well suited for studies of development and neurobiology, typically examined in young animals (Brenner 1974). This included 50 mM supplemental NaCl added to standard medium. Because *C. elegans* is now a prominent model for studies of aging, it is important to understand how culture conditions affect aging and lifespan. Here we demonstrate that 50 mM supplemental NaCl added to standard medium indeed affects aging. Animals cultured in reduced NaCl medium displayed an extended lifespan and delayed age-related degeneration of body movement and pharyngeal pumping. The extended lifespan caused by these culture conditions was also observed in three long-lived mutants: *daf-2* and *age-1,* which have reduced insulin/IGF-1 signaling, and *nuo-6,* which has reduced mitochondrial activity. These findings suggest that the mechanism of lifespan extension caused by reduced NaCl may be distinct from mechanisms of lifespan extension caused by decreased insulin/IGF-1 signaling or mitochondrial activity. However, it is also possible that the lifespan extension effects are additive because all these interventions are only partial; the *daf-2* and *age-1* alleles we analyzed cause a partial loss-of-function, and the *nuo-6* mutants have reduced - but not abolished - mitochondrial activity (Yang and Hekimi 2010b). Reduced NaCl medium did not extend the lifespan of the short-lived mutants *daf-16* and *daf-12*, nor did it further extend the lifespans of *eat-2* and *che-1* mutants. Our results support the model that 50 mM NaCl in standard medium is tolerated in young animals but mildly toxic in old adults, since it accelerates aging and reduces lifespan. Three other aspects of standard culture conditions have been demonstrated to influence lifespan. (1) Reducing temperature to 15°C extends lifespan, whereas increasing temperature to 25°C shortens lifespan (Klass 1977; Huang et al. 2004). (2) Reducing the pathogenicity of *E. coli* OP50 extends lifespan; this can be accomplished with antibiotic treatment of *E. coli* OP50 (Garigan et al. 2002) or by substituting a non-pathogenic bacterial food source (Yu et al. 2015). (3) Higher oxygen levels (60% and 80%) reduce lifespan, whereas hypoxia (1% oxygen) extends lifespan (Adachi et al. 1998).

*C. elegans* has evolved sophisticated mechanisms to respond to osmotic stress, and a central strategy is to synthesize the small osmolyte glycerol, which helps animals retain water (Urso and Lamitina 2021). Despite these defenses, our results show that excess NaCl caused multiple toxicities: it delayed the rate of development from egg to sexual maturity, reduced progeny production, accelerated the age-related declines of neuromuscular activities including body movement and pharyngeal pumping, and reduced mean lifespan in both wild-type and mutant animals (*daf-2, age-1, daf-16, daf-12, nuo-6, eat-2,* and *che-1*). These results are consistent with previous studies showing that excess NaCl can reduce nematode survival (Lamitina et al. 2004; Dmitrieva and Burg 2007; Chandler-Brown et al. 2015; Anderson et al. 2016). The *gpdh-1* gene promotes synthesis of glycerol, and transcription is activated by excess NaCl (Urso and Lamitina 2021). We confirmed this observation in young adults. Furthermore, we investigated *gpdh-1* levels throughout adulthood. Unexpectedly, the gene is not activated in middle aged adults, but activation resumes in older adults. Thus, transcriptional activation of *gpdh-1* requires excess NaCl but is also sensitive to adult age. We speculate that activation of the *gpdh-1* in young adults may result in sufficient glycerol accumulation to restore homeostasis, and this effect persists into middle aged adults, resulting in *gpdh-1* mRNA levels returning to baseline. Older adults may eventually lose this glycerol and/or be more susceptible to external NaCl, and thus resume activation of *gpdh-1* transcription. Animals cultured on reduced NaCl and standard NGM expressed *gpdh-1* to the same extent, indicating that the concentration of NaCl in standard NGM is not sufficient to upregulate *gpdh-1* transcription. We speculate that 50 mM supplemental NaCl in standard NGM may reduce lifespan because it is toxic, but is not sufficient to induce an osmotic stress response.

Older adults exposed to excess NaCl also induce expression of *hsp-6*, indicating that excess NaCl may also stimulate the mitochondrial unfolded protein response. In young adults exposed to excess NaCl, expression of *hsp-4* and *gst-4* are reduced, suggesting these pathways may be inhibited by excess NaCl. Notably, glutathione S-transferase has been shown to be relevant for salt tolerance in plants (Hao et al. 2021; Gao et al. 2022). In day 0 adults, standard NGM reduced the expression of *gst-4*, a gene involved in drug metabolism, compared to reduced NaCl. It may be that higher expression of *gst-4* contributes to the health of animals cultured in reduced NaCl. Our results have practical implications for conducting and interpreting lifespan and aging experiments with *C. elegans*. We recommend that NGM dishes used for lifespan experiments not be stored for long time periods, because evaporative loss can increase the concentration of NaCl and thus alter the experimental outcome. When investigating lifespan-extending drugs or mutations, it may be valuable to investigate them in NGM dishes with 0 mM supplemental NaCl as well, since this will illuminate how the intervention interacts with NaCl toxicity.

## Materials and Methods

### General maintenance and NGM preparation

All *C. elegans* strains were cultured using standard methods (Brenner 1974) unless otherwise stated. Standard NGM dishes contain 3.0 g of supplemental NaCl per liter, which results in 51.3 mM supplemental NaCl. For simplicity, we refer to standard NGM as having 50 mM supplemental NaCl. **Table S1** shows the formula used to prepare reduced and excess NaCl dishes. *C. elegans* were maintained on NGM dishes seeded with 200 μL *Escherichia coli* strain OP50 at 20°C in the dark. Strains were provided by the *Caenorhabditis* Genetic Center (CGC), University of Minnesota. **Table S2** provides a list of all strains.

**Table S1.** Preparation of medium.

| Supplemental NaCl (mM) (actual) | Supplemental NaCl (mM) (name in paper) | Agar (g/L) | Peptone (g/L) | NaCl (g/L) |
| --- | --- | --- | --- | --- |
| 0 | 0 (reduced NaCl medium) | 17.0 | 2.5 | 0.0 |
| 51.3 | 50 (standard medium) | 17.0 | 2.5 | 3.0 |
| 200 | 200 (excess NaCl medium) | 17.0 | 2.5 | 11.68 |

**Table S2.**
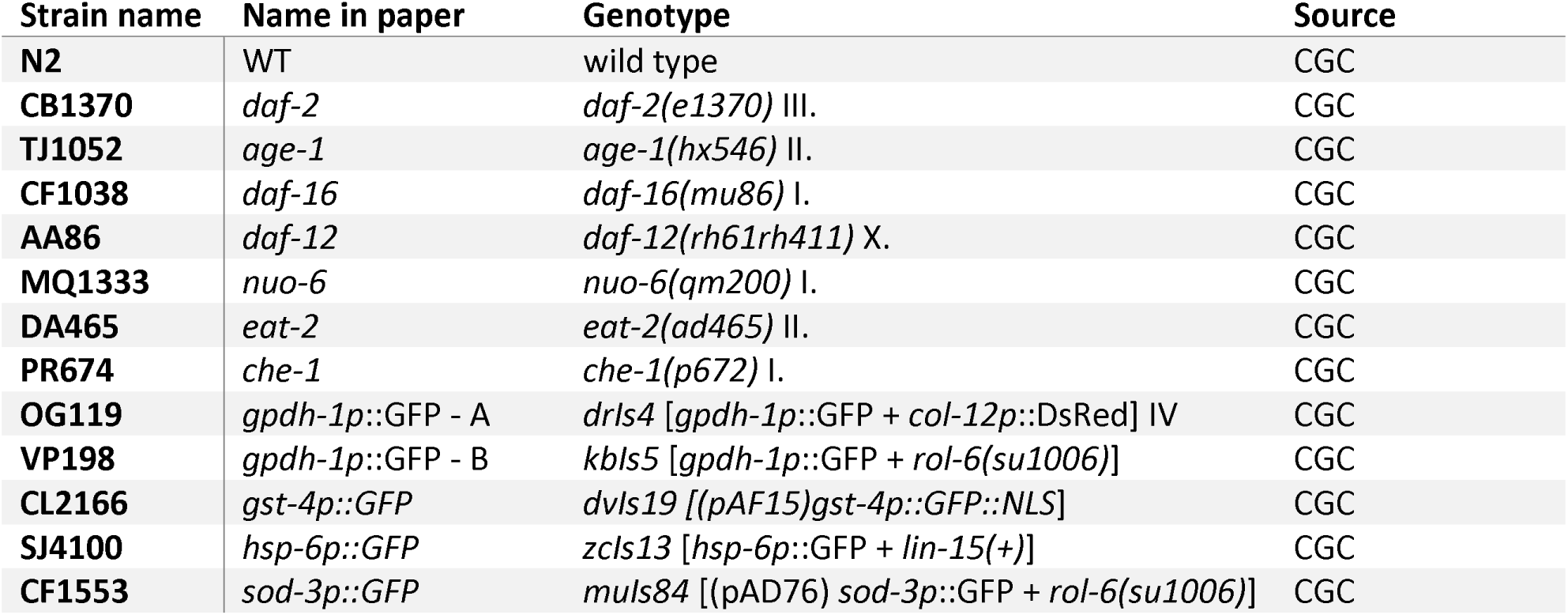
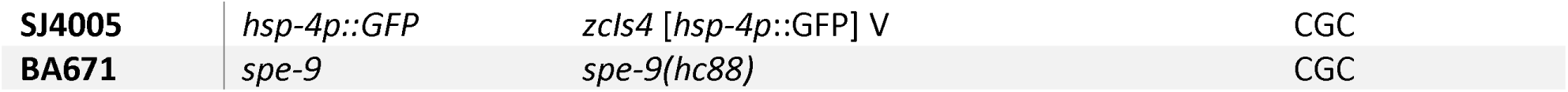
*C. elegans* strains.

| Strain name | Name in paper | Genotype | Source |
| --- | --- | --- | --- |
| N2 | WT | wild type | CGC |
| CB1370 | *daf-2* | *daf-2(e1370)* III. | CGC |
| TJ1052 | *age-1* | *age-1(hx546)* II. | CGC |
| CF1038 | *daf-16* | *daf-16(mu86)* I. | CGC |
| AA86 | *daf-12* | *daf-12(rh61rh411)* X. | CGC |
| MQ1333 | *nuo-6* | *nuo-6(qm200)* I. | CGC |
| DA465 | *eat-2* | *eat-2(ad465)* II. | CGC |
| PR674 | *che-1* | *che-1(p672)* I. | CGC |
| OG119 | *gpdh-1p::GFP* - A | *drls4* [*gpdh-1p::GFP* + *col-12p::DsRed*] IV | CGC |
| VP198 | *gpdh-1p::GFP* - B | *kbls5* [*gpdh-1p::GFP* + *rol-6(su1006)*] | CGC |
| CL2166 | *gst-4p::GFP* | *dvls19* [(*pAF15*)*gst-4p::GFP::NLS*] | CGC |
| SJ4100 | *hsp-6p::GFP* | *zcls13* [*hsp-6p::GFP* + *lin-15(+)*] | CGC |
| CF1553 | *sod-3p::GFP* | *muls84* [(*pAD76*) *sod-3p::GFP* + *rol-6(su1006)*] | CGC |
| **SJ4005** | *hsp-4p::GFP* | *zcls4 [hsp-4p::GFP] V* | CGC |
| **BA671** | *spe-9* | *spe-9(hc88)* | CGC |

To synchronize animals for NaCl supplementation experiments, we cultured 5 adult worms on standard NGM dishes for 4 days to generate a mixed population of worms, then transferred adults from that dish to NGM dishes with the appropriate NaCl concentration. On excess NaCl, animals have a lower rate of egg laying; to obtain an equal number of eggs as those grown on reduced NaCl or standard NGM, we used more adults and allowed them to lay eggs for a longer period of time. We transferred 10 adults/dish for reduced NaCl medium and standard medium, and 20 adults/dish for excess NaCl medium. These adults were allowed to lay eggs for 1-2 hours for reduced NaCl medium and standard medium, and 2-4 hours for excess NaCl medium. We then removed adult worms, thus generating a population of age-synchronized eggs (**Fig.S2A**). Experiments started 3 days (72 hours) later with a synchronized cohort of day 0 adults.

To evaluate evaporative loss from NGM dishes, we pre-weighed empty Petri dishes (60 mm), dispensed into them 10 mL of molten NGM media, dried them at room temperature for 1-2 days, seeded them with 250 µL *E. coli* OP50, and dried them at room temperature for an additional 2 days. Dishes were then weighed, and the weight of the empty dish was subtracted to determine the weight of NGM at day 0. Dishes were stored at 4°C or 20°C and weighed every 7 days or every 2-3 days, respectively, to monitor evaporative weight loss.

### Inductively coupled plasma mass spectrometry (ICP-MS)

We performed elemental analysis on agar (A7002-1KG, Sigma-Aldrich) and peptone (Bacto™Peptone, 211677 Gibco, ordered through Thermo Fisher Scientific), which are ingredients of NGM, and yeast extract (Bacto™ Yeast Extract, 212750, Gibco, ordered through Thermo Fisher Scientific) and tryptone (Bacto™ Trypton, 211705, Gibco), which are ingredients of LB. For each ingredient, we performed analysis on three samples, each from the same vendor.

Elemental analysis was performed by the Northwestern University Quantitative Bio-element Imaging Center. Samples were digested in 500 µL concentrated trace-grade nitric acid (>69%, Thermo Fisher Scientific, Waltham, MA, USA) and 125 µL trace-grade hydrogen peroxide (>30%, GFS Chemicals, Columbus, OH, USA) at 65°C for at least 3 hours to allow for complete sample digestion. Ultra-pure H_2_O (18.2 MΩ·cm) was added to produce a final solution of 5.0% nitric acid in a total sample volume of 10 mL. We measured sodium (Na), arsenic (As), calcium (Ca), cadmium (Cd), cobalt (Co), chromium (Cr), copper (Cu), iron (Fe), potassium (K), magnesium (Mg), manganese (Mn), nickel (Ni), selenium (Se), vanadium (V), and zinc (Zn). Quantitative standards were made using a mixed elemental standard (Inorganic Ventures, Christiansburg, VA, USA) containing 100 µg/mL each of Na, As, Ca, Cd, Co, Cr, Cu, Fe, K, Mg, Mn, Ni, Se, V, and Zn. This was diluted to create (1) a 100 ng/g mixed element standard in 5.0% nitric acid (v/v) in a total sample volume of 50 mL and (2) a 10 ng/g mixed element standard in 5.0% nitric acid (v/v) in a total sample volume of 50 mL. A 1000 µg/mL stock standard containing Na, Mg, K, and Ca (IV-STOCK-3 from Inorganic Ventures) was diluted to create a 25 µg/g mixed element standard in 5.0% nitric acid. A solution of 5.0% nitric acid (v/v) was used as the calibration blank.

ICP-MS was performed on a computer-controlled (QTEGRA software) Thermo iCapQ ICP-MS (Thermo Fisher Scientific, Waltham, MA, USA) operating in KED mode and equipped with an ESI SC-2DX PrepFAST autosampler (Omaha, NE, USA) as previously described (Mendoza et al. 2017). Internal standard was added inline using the prepFAST system and consisted of 1 ng/mL of a mixed element solution containing Bi, In, ^6^Li, Sc, Tb, Y (IV-ICPMS-71D from Inorganic Ventures). Online dilution was carried out by the prepFAST system and used to generate a calibration curve consisting of 100, 50, 20, 10, 5, 2, 1, 0.2, 0.1, and 0.05 parts per billion (ppb) of all elements. Additionally, online dilution was used to further calibrate Na, Mg, K and Ca at 25, 12.5, 5, 2, 1, 0.5, and 0.25 parts per million (ppm). Each sample was acquired using 1 survey run (10 sweeps) and 3 main (peak jumping) runs (40 sweeps). The isotopes selected for analysis were ^23^Na, ^24^Mg, ^39^K, ^44^Ca, ^51^V, ^52^Cr, ^55^Mn, ^56^Fe, ^57^Fe, ^59^Co, ^60^Ni, ^62^Ni, ^63^Cu, ^65^Cu, ^66^Zn, ^68^Zn, ^75^As, ^77^Se, ^111^Cd, and ^45^Sc, ^89^Y, ^115^In (chosen as internal standards for data interpolation and machine stability). Instrument performance was optimized daily through autotuning followed by verification via a performance report (passing manufacturer specifications).

### Lifespan

To measure mean lifespan, we cultured hermaphrodites beginning at the embryo stage on NGM dishes containing 0, 50, or 200 mM supplemental NaCl and seeded with 250 μL *E. coli* OP50. Beginning 72 hours later (day 0 adults), survival was monitored daily. Hermaphrodites were transferred to new dishes daily during the reproductive period to remove self-progeny, and every 2-3 days thereafter. Hermaphrodites that died due to matricidal hatching, gonad extrusion, or desiccation on the edge of the dish were scored as alive on the day prior to death and were censored from the experiment for each day thereafter. Animals were determined to be dead by observation with a dissecting microscope and gentle manipulation with a sterile platinum wire; if no spontaneous or induced movement was observed, they were scored as dead. Experiments continued until all animals had died or been censored. The data were analyzed by Kaplan-Meier Analysis using GraphPad Prism Version 10.2.2 software, with statistical significance considered at p<0.05 using the log-rank (Mantel-Cox) test. Statistical significance was adjusted for comparison of 3 curves based on the Bonferroni method. The family-wise significance level (0.05, 0.01, and 0.001) was divided by K, the count of the number of comparisons, i.e. 3. Hence adjusted significance levels are: p<0.00833*, p<0.00166**, and p<0.000166***.

### Measurement of body movement and pharyngeal pumping rate

Wild-type animals were cultured as described for lifespan experiments and observed daily throughout their adult lifespan with a dissecting microscope. Body movement was measured by picking animals individually onto NGM dishes containing 0, 50, or 200 mM supplemental NaCl. The number of sinusoidal body movements was counted during a 15 second interval, and that number was multiplied by four to calculate body movements per minute. One sinusoidal body movement was defined as the movement of the animal’s head from its left-most position to its right-most position or *vice versa*, while the animal was moving either forward or backward. In addition, for two of the body movement experiments we also categorized animals as one of three possible movement classes using the following rules: class A - vigorous, coordinated, spontaneous movement; class B - uncoordinated movement: part of body (head, tail, etc.) is paralyzed or uncoordinated while the rest of the body can move normally; class C - majority of the animal is paralyzed, no forward or backward movement, movement is only observed in head or tail (Newell Stamper et al. 2018). In this experiment the same animals were observed at multiple time points; however, individual animals were not separately tracked. The same animals were used to measure the pharyngeal pumping rate by using a dissecting microscope to count the number of pharynx pumps during a 15 second interval.

### Time-series RNA-seq

Information on the RNA-seq experiment may be found in previous publications (Egan et al. 2024; Mosley et al. 2024); the same dataset was used for the experiments presented here.

### Measurement of development rate, daily self-progeny, and total self-progeny

Wild-type animals were cultured as described for lifespan experiments. To measure development rate, we placed animals as eggs onto individual NGM dishes and began monitoring them ∼66 hours later. Each hour thereafter, we examined each animal for the presence of eggs on the dish. For each individual animal, we defined the ‘time to first progeny’ as the number of hours elapsed between when they were placed on the dish and when they laid their first egg.

Daily self-progeny were measured starting on day 0 of adulthood. We placed individual hermaphrodites on a dish with 0, 50, or 200 mM supplemental NaCl. Each day, we transferred the adults onto fresh dishes and retained the dishes bearing their progeny. Dishes with progeny were cultured at 15°C or 20°C and, 2-4 days later, live progeny were counted then discarded using a glass pipette and a vacuum pump. These longitudinal data directly measure daily progeny number and were used to calculate the total number of progeny (day 0 to day 5 of adulthood).

### Assessment of autofluorescence and stress response induction using reporter strains for *gpdh-1*, *gst-4*, *sod-3, hsp-4,* and *hsp-6*

Transgenic animals containing one reporter construct were cultured as described for lifespan experiments. Starting with a synchronized batch of eggs, animals were cultured for 72±4 hours. To analyze worms by fluorescence microscopy, we prepared 3% agar slides and added a drop of levamisol (3 mM, in M9) onto the agar as previously described (Pohl et al. 2023). If possible ∼10 worms were added per slide. As necessary, worms were oriented using an eye lash. Excess levamisol was removed if needed, the worms were covered with a cover slide, and the cover slide was sealed with 3% agar. Brightfield and fluorescence images (excitation (ex.) λ=475 nm; emission (em.) λ=510 nm) of animals were acquired using a Leica DMi8 microscope. Fluorescence intensity of each worm was measured using Fiji (ImageJ, 1.53f51). The experiments were repeated at least three times independently unless otherwise indicated. For the purpose of showing representative images, the same level of brightness and contrast adjustment was applied to all images; however, quantitative analysis was conducted on raw images without any adjustments. To analyze days 1-16 of adulthood, we transferred adult hermaphrodites to new dishes daily during their reproductive period, to separate them from their progeny, and then transferred them to new dishes every 2-3 days thereafter. Autofluorescence in WT animals was measured on day 0, 5, and 11 of adulthood at the following channels: (i) red channel (ex. λ=555 nm; em. λ=590 nm), (ii) green channel (ex. λ=475 nm; em. λ=510 nm), and (iii) blue channel (ex. λ=390 nm; em. λ=435 nm).

### Statistics

All statistical analyses were performed in GraphPad Prism, Version 10.2.3 (403). Experiments were analyzed using one-way or two-way ANOVA with either Dunnette’s, Bonferroni’s or Tukey’s multiple comparison analysis or mixed-effects model with Geisser-Greenhouse correction, as indicated in the Figure legends. *, **, and *** indicate statistical significance with p<0.05, p<0.01, and p<0.001, respectively. For lifespan experiments, we used Kaplan-Meier Analysis with log-rank test for trend; p-values were corrected using Bonferroni correction, where the Bonferroni-corrected threshold α_Bonferroni_ = α_family_/K, K=number of comparisons i.e. 3, and α_family_=0.05, 0.01, or 0.001 for *, **, or ***, respectively. If the p-value was lower than the Bonferroni-corrected threshold, i.e. 0.0167, 0.0034, pr 0.00034, the respective significant level, i.e. *, ** or ***, was applied. For graphs with many comparisons, such as Figure 3, we used the compact letter display for comparison; for any two groups that share a letter, p>0.05 and no significant difference was observed; for groups that do not share a letter, p<0.05 and a statistically significant difference was observed.

---

**Supplementary Figure 1.**
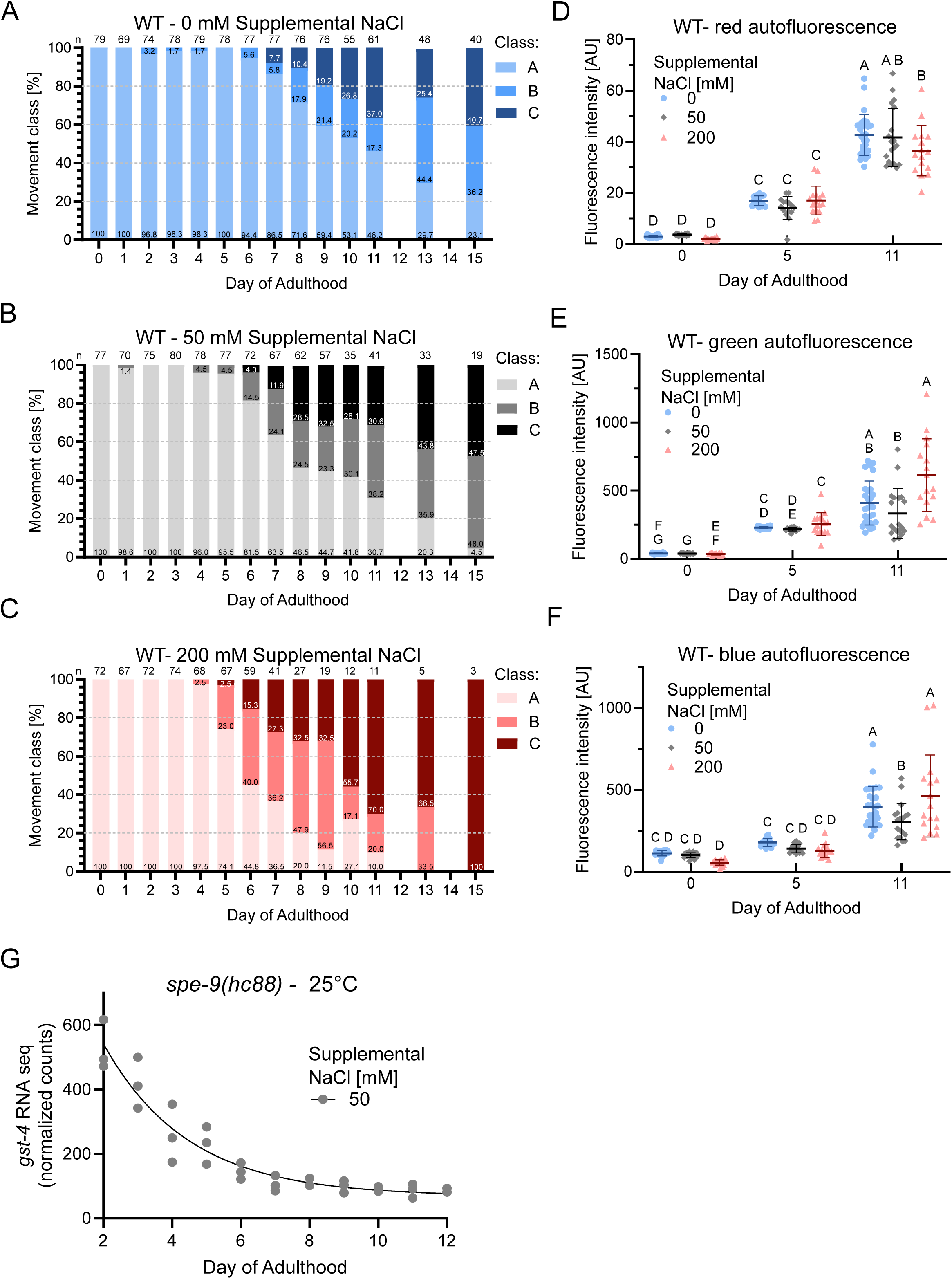
Effect of supplemental NaCl on age-related changes of body movement and autofluorescence, and age-related changes of *gst-4* RNA expression levels. **A-C)** Wild-type hermaphrodites were categorized into movement classes on adult days 0-15 using a dissecting microscope. Class A: vigorous, coordinated, spontaneous movement; Class B: uncoordinated movement - part of body (head, tail, etc.) is paralyzed or uncoordinated; Class C: majority of animal is paralyzed, no forward or backward movement, only moves head or tail slightly. Bars are color coded and labeled with the percent of animals in each class, which sum to 100%. Two independent experiments (number of animals (n) shown above each day). Day 8 data are shown in Figure 1D. **D-F)** Autofluorescence displayed by individual wild-type animals was measured on day 0, 5, and 11 of adulthood using the following channels: **D)** red channel (excitation λ=555 nm; emission λ=590 nm). **E)** green channel (excitation λ=475 nm; emission λ=510 nm). **F)** blue channel (excitation λ=390 nm; emission λ=435 nm). Mean±SD. Two-way ANOVA with Tukey’s multiple comparisons test was performed using the compact letter display for comparison: for any two groups that share a letter, p>0.05, indicating no significant difference; for groups with different letters, p<0.05, indicating a statistically significant difference. Two independent experiments with n≥6. **G)** *gst-4* RNA expression levels were measured by RNA-seq in *spe-9(hc88)* mutant animals cultured at 25°C on standard NGM dishes (50 mM supplemental NaCl). The *spe-9* mutation prevents progeny production at 25°C. Three biological replicates per day.

**Supplementary Figure 2.**
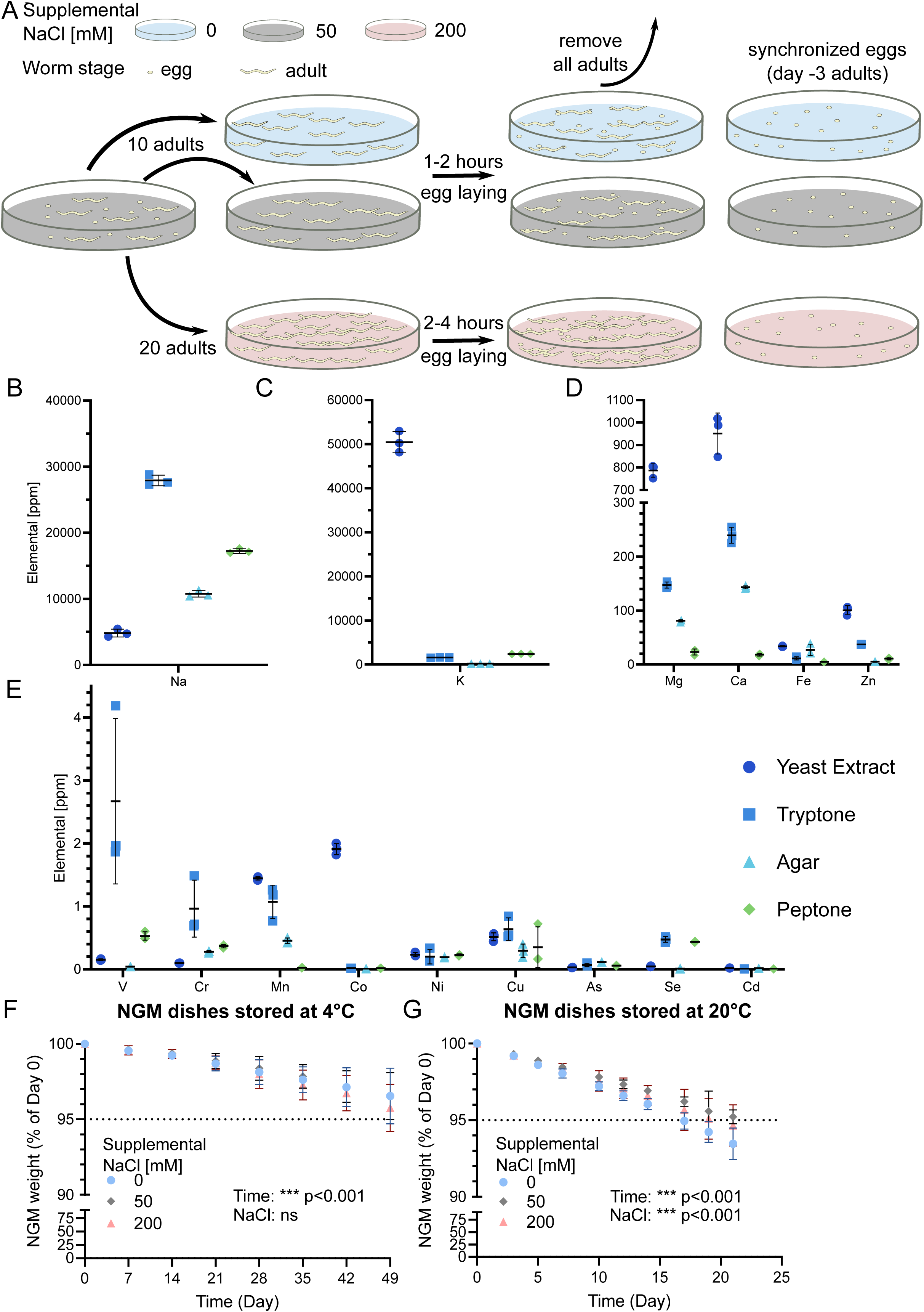
Procedure for synchronizing eggs, elemental analysis of NGM and LB ingredients, and measurement of NGM dish evaporative loss. **A)** Diagrams illustrate the experimental method used to obtain synchronized populations of eggs. Hermaphrodites were initially cultured on standard NGM containing 50 mM supplemental NaCl, then were transferred to NGM dishes containing 0, 50, or 200 mM supplemental NaCl. Adults were allowed to briefly lay eggs and then were removed, and the dish with synchronized eggs was retained. For 0 and 50 mM supplemental NaCl medium, we used 10 adults for 1-2 hours to obtain a sufficient number of eggs (designated day −3 adults). For 200 mM supplemental NaCl medium, we used 20 adults for 2-4 hours to obtain a similar number of eggs, since this concentration of NaCl has the toxic effect of reducing egg laying. **B-E)** Using ICP-MS, we analyzed the dry ingredients that are used to formulate NGM and LB: yeast extract, tryptone, agar and peptone. Data points are three independent replicates in parts per million (ppm), and bars are mean±S.D. Panel B is sodium (Na), panel C is potassium (K), panel D is magnesium (Mg), calcium (Ca), iron (Fe), and zinc (Zn), and panel E is vanadium (V), chromium (Cr), manganese (Mn), cobalt (Co), nickel (Ni), copper (Cu), arsenic (As), selenium (Se), and cadmium (Cd). Note that each panel has different vertical axis values. **F-G)** Petri dishes containing NGM medium were weighed periodically, and the weight of the medium was calculated by subtracting the weight of the empty dish. Dishes were stored at 4°C or 20°C. Values are mean±S.D. Three independent experiments with a combined n=30 for each condition at day 0. Contaminated plates were removed leading to n=27, 25, and 27 at the end of study for 0, 50 and 200 mM NaCl, respectively. Mixed-effects model with Geisser-Greenhouse correction for comparison between time, NaCl, and any interactions. Dishes at 4°C displayed a time-dependent loss of weight that was less than 5% after 49 days, and this was not significantly affected by the NaCl concentration. Dishes at 20°C displayed a time-dependent loss of weight that was more rapid than at 4°C, and NaCl concentration showed a significant effect on evaporation, which was highest for 0 mM supplemental NaCl.

## Data availability

Strains are available upon request. The authors affirm that all data necessary for confirming the conclusions of the article are present within the article, figures, and tables.

## Author Contributions

Conceptualization: FP, BME, KK; Data Curation: FP, BME, MCM; Formal Analysis: FP, MCM; Funding Acquisition: FP, KK; Investigation: FP, BME, DLS, MCM, MAG, SH, CHC; Methodology: FP, BME, DLS, KK; Project Administration: FP, BME, DLS, KK; Supervision: FP, BME, DLS, KK; Software: MCM; Validation: FP, BME, KK; Visualization: FP; Writing, Original Draft Preparation: FP, BME; Writing, Review and Editing: FP, BME, KK

## Acknowledgements

We thank the *Caenorhabditis* Genetics Center (CGC; https://cgc.umn.edu/), which is funded by the National Institutes of Health Office of Research Infrastructure Programs (P40 OD010440), for mutant nematode strains. Elemental analysis was performed at the Northwestern University Quantitative Bio-element Imaging Center generously supported by the NIH under Grant S10OD020118, and we thank Rebecca Sponenburg for conducting this analysis.

## References

Adachi H, Fujiwara Y, Ishii N. 1998. Effects of Oxygen on Protein Carbonyl and Aging in Caenorhabditis elegans Mutants With Long (age-1) and Short (mev-1) Life Spans. The Journals of Gerontology: Series A. 53A(4):B240–B244. doi:10.1093/GERONA/53A.4.B240. [accessed 2024 Sep 19]. 10.1093/gerona/53A.4.B240.

Al-Amin M, Kawasaki I, Gong J, Shim YH. 2016. Caffeine Induces the Stress Response and Up-Regulates Heat Shock Proteins in Caenorhabditis elegans. Mol Cells. 39(2):163–168. doi:10.14348/MOLCELLS.2016.2298.

Anderson EN, Corkins ME, Li JC, Singh K, Parsons S, Tucey TM, Sorkaç A, Huang H, Dimitriadi M, Sinclair DA, et al. 2016. C. elegans lifespan extension by osmotic stress requires FUdR, base excision repair, FOXO, and sirtuins. Mech Ageing Dev. 154:30–42. doi:10.1016/J.MAD.2016.01.004.

Bar-Ziv R, Bolas T, Dillin A. 2020. Systemic effects of mitochondrial stress. EMBO Rep. 21(6). doi:10.15252/embr.202050094. [accessed 2020 Aug 30]. https://www.embopress.org/doi/10.15252/embr.202050094.

Bar-Ziv R, Frakes AE, Higuchi-Sanabria R, Bolas T, Frankino PA, Gildea HK, Metcalf MG, Dillin A. 2020. Measurements of physiological stress responses in C. Elegans. Journal of Visualized Experiments. 2020(159):1–21. doi:10.3791/61001. [accessed 2021 Mar 4]. www.jove.comURL:https://www.jove.com/video/61001.

Berendzen KM, Durieux J, Shao L-W, Tian Y, Kim H, Wolff S, Liu Y, Dillin A. 2016. Neuroendocrine Coordination of Mitochondrial Stress Signaling and Proteostasis. Cell. 166(6):1553–1563.e10. doi:10.1016/j.cell.2016.08.042. [accessed 2020 Feb 24]. https://linkinghub.elsevier.com/retrieve/pii/S0092867416311394.

Bourque CW. 2008. Central mechanisms of osmosensation and systemic osmoregulation. Nature Reviews Neuroscience 2008 9:7. 9(7):519–531. doi:10.1038/nrn2400. [accessed 2022 Oct 31]. https://www.nature.com/articles/nrn2400.

Brenner S. 1974. The genetics of Caenorhabditis elegans. Genetics. 77(1):71–94. doi:10.1093/GENETICS/77.1.71. [accessed 2023 Jul 23]. https://pubmed.ncbi.nlm.nih.gov/4366476/.

Chandler-Brown D, Choi H, Park S, Ocampo BR, Chen S, Le A, Sutphin GL, Shamieh LS, Smith ED, Kaeberlein M. 2015. Sorbitol treatment extends lifespan and induces the osmotic stress response in Caenorhabditis elegans. Front Genet. 6(OCT):160335. doi:10.3389/FGENE.2015.00316/BIBTEX. [accessed 2024 Aug 9]. www.frontiersin.org.

Choe KP. 2013. Physiological and molecular mechanisms of salt and water homeostasis in the nematode Caenorhabditis elegans. Am J Physiol Regul Integr Comp Physiol. 305(3). doi:10.1152/AJPREGU.00109.2013. [accessed 2024 Apr 8]. https://pubmed.ncbi.nlm.nih.gov/23739341/.

Dillin A, Hsu AL, Arantes-Oliveira N, Lehrer-Graiwer J, Hsin H, Fraser AG, Kamath RS, Ahringer J, Kenyon C. 2002. Rates of behavior and aging specified by mitochondrial function during development. Science (1979). 298(5602):2398–2401. doi:10.1126/SCIENCE.1077780/SUPPL_FILE/DILLIN.SOM.REV.PDF. [accessed 2024 Sep 17]. https://www.science.org/doi/10.1126/science.1077780.

Dmitrieva NI, Burg MB. 2007. High NaCl Promotes Cellular Senescence. Cell Cycle. 6(24):3108–3113. doi:10.4161/CC.6.24.5084. [accessed 2025 Feb 27]. https://www.tandfonline.com/doi/abs/10.4161/cc.6.24.5084.

Dues DJ, Schaar CE, Johnson BK, Bowman MJ, Winn ME, Senchuk MM, Van Raamsdonk JM. 2017. Uncoupling of Oxidative Stress Resistance and Lifespan in Long-lived isp-1 Mitochondrial Mutants in Caenorhabditis elegans. Free Radic Biol Med. 108:362. doi:10.1016/J.FREERADBIOMED.2017.04.004. [accessed 2024 Sep 17]. /pmc/articles/PMC5493208/.

Egan BM, Pohl F, Anderson X, Williams SC, Adodo IG, Hunt P, Wang Z, Chiu CH, Scharf A, Mosley M, et al. 2024. The ACE inhibitor captopril inhibits ACN-1 to control dauer formation and aging. Development (Cambridge). 151(3):1–16. doi:10.1242/DEV.202146/342404/AM/THE-ACE-INHIBITOR-DRUG-CAPTOPRIL-INHIBITS-ACN-1-TO. [accessed 2024 Sep 17]. 10.1242/dev.202146.

Ewbank JJ, Barnes TM, Lakowski B, Lussier M, Bussey H, Hekimi S. 1997. Structural and Functional Conservation of the Caenorhabditis elegans Timing Gene clk-1. Science (1979). 275(5302):980–983. doi:10.1126/SCIENCE.275.5302.980. [accessed 2024 Sep 17]. https://www.science.org/doi/10.1126/science.275.5302.980.

Félix MA, Braendle C. 2010. The natural history of Caenorhabditis elegans. Current Biology. 20(22):R965–R969. doi:10.1016/J.CUB.2010.09.050.

Félix MA, Duveau F. 2012. Population dynamics and habitat sharing of natural populations of Caenorhabditis elegans and C. briggsae. BMC Biol. 10(1):1–19. doi:10.1186/1741-7007-10-59/FIGURES/6. [accessed 2024 Sep 19]. https://bmcbiol.biomedcentral.com/articles/10.1186/1741-7007-10-59.

Felkai S, Ewbank JJ, Lemieux J, Labbé JC, Brown GG, Hekimi S. 1999. CLK-1 controls respiration, behavior and aging in the nematode Caenorhabditis elegans. EMBO J. 18(7):1783–1792. doi:10.1093/EMBOJ/18.7.1783. [accessed 2024 Sep 17]. https://www.embopress.org/doi/10.1093/emboj/18.7.1783.

Feng J, Bussière F, Hekimi S. 2001. Mitochondrial Electron Transport Is a Key Determinant of Life Span in Caenorhabditis elegans. Dev Cell. 1(5):633–644. doi:10.1016/S1534-5807(01)00071-5.

Ferguson GD, Bridge WJ. 2019. The glutathione system and the related thiol network in Caenorhabditis elegans. Redox Biol. 24:101171. doi:10.1016/j.redox.2019.101171. [accessed 2020 Jan 28]. http://www.ncbi.nlm.nih.gov/pubmed/30901603.

Friedman DB, Johnson TE. 1988. A mutation in the age-1 gene in Caenorhabditis elegans lengthens life and reduces hermaphrodite fertility. Genetics. 118(1):75–86. doi:10.1093/GENETICS/118.1.75. [accessed 2024 Sep 16]. https://pubmed.ncbi.nlm.nih.gov/8608934/.

Gao H, Yu C, Liu R, Li Xiaoyan, Huang H, Wang X, Zhang C, Jiang N, Li Xiaofang, Cheng S, et al. 2022. The Glutathione S-Transferase PtGSTF1 Improves Biomass Production and Salt Tolerance through Regulating Xylem Cell Proliferation, Ion Homeostasis and Reactive Oxygen Species Scavenging in Poplar. Int J Mol Sci. 23(19). doi:10.3390/IJMS231911288/S1. [accessed 2024 Aug 9]. /pmc/articles/PMC9569880/.

Garigan D, Hsu AL, Fraser AG, Kamath RS, Abringet J, Kenyon C. 2002. Genetic analysis of tissue aging in Caenorhabditis elegans: a role for heat-shock factor and bacterial proliferation. Genetics. 161(July):1101–1112.

Guisnet A, Maitra M, Pradhan S, Hendricks M. 2021. A three-dimensional habitat for C. elegans environmental enrichment. PLoS One. 16(1). doi:10.1371/JOURNAL.PONE.0245139. [accessed 2024 Sep 19]. /pmc/articles/PMC7799825/.

Hao Y, Xu S, Lyu Z, Wang H, Kong L, Sun S. 2021. Comparative Analysis of the Glutathione S-Transferase Gene Family of Four Triticeae Species and Transcriptome Analysis of GST Genes in Common Wheat Responding to Salt Stress. Int J Genomics. 2021(1):6289174. doi:10.1155/2021/6289174. [accessed 2024 Aug 9]. https://onlinelibrary.wiley.com/doi/full/10.1155/2021/6289174.

Huang CG, Lamitina T, Agre P, Strange K. 2007. Functional analysis of the aquaporin gene family in Caenorhabditis elegans. Am J Physiol Cell Physiol. 292(5). doi:10.1152/AJPCELL.00514.2006. [accessed 2024 Sep 19]. https://pubmed.ncbi.nlm.nih.gov/17229810/.

Huang C, Xiong C, Kornfeld K. 2004. Measurements of age-related changes of physiological processes that predict lifespan of Caenorhabditis elegans. Proc Natl Acad Sci U S A. 101(21):8084–8089. doi:10.1073/PNAS.0400848101/ASSET/8E30DB6E-62F5-440F-BD9A-53D481A4513F/ASSETS/GRAPHIC/ZPQ0210449330004.JPEG. [accessed 2022 Mar 15]. www.pnas.orgcgidoi10.1073pnas.0400848101.

Kenyon C, Chang J, Gensch E, Rudner A, Tabtiang R. 1993. A C. elegans mutant that lives twice as long as wild type. Nature 1993 366:6454. 366(6454):461–464. doi:10.1038/366461a0. [accessed 2024 Sep 16]. https://www.nature.com/articles/366461a0.

Kimura KD, Tissenbaum HA, Liu Y, Ruvkun G. 1997. daf-2, an insulin receptor-like gene that regulates longevity and diapause in Caenorhabditis elegans. Science. 277(5328):942–946. doi:10.1126/SCIENCE.277.5328.942. [accessed 2024 Sep 16]. https://pubmed.ncbi.nlm.nih.gov/9252323/.

Kingsley SF, Seo Y, Allen C, Ghanta KS, Finkel S, Tissenbaum HA. 2021. Bacterial processing of glucose modulates C. elegans lifespan and healthspan. Scientific Reports 2021 11:1. 11(1):1–12. doi:10.1038/s41598-021-85046-3. [accessed 2024 Sep 17]. https://www.nature.com/articles/s41598-021-85046-3.

Kitisin T, Muangkaew W, Sukphopetch P. 2022. Caenorhabditis elegans DAF-16 regulates lifespan and immune responses to Cryptococcus neoformans and Cryptococcus gattii infections. BMC Microbiol. 22(1):1–12. doi:10.1186/S12866-022-02579-X/FIGURES/6. [accessed 2024 Sep 17]. https://bmcmicrobiol.biomedcentral.com/articles/10.1186/s12866-022-02579-x.

Klass MR. 1977. Aging in the nematode Caenorhabditis elegans: Major biological and environmental factors influencing life span. Mech Ageing Dev. 6(C):413–429. doi:10.1016/0047-6374(77)90043-4.

Komura T, Yamanaka M, Nishimura K, Hara K, Nishikawa Y. 2021. Autofluorescence as a noninvasive biomarker of senescence and advanced glycation end products in Caenorhabditis elegans. npj Aging and Mechanisms of Disease 2021 7:1. 7(1):1–11. doi:10.1038/s41514-021-00061-y. [accessed 2024 Sep 17]. https://www.nature.com/articles/s41514-021-00061-y.

Kumar S, Egan BM, Kocsisova Z, Schneider DL, Murphy JT, Diwan A, Kornfeld K. 2019. Lifespan Extension in C. elegans Caused by Bacterial Colonization of the Intestine and Subsequent Activation of an Innate Immune Response. Dev Cell. 49(1):100–117.e6. doi:10.1016/j.devcel.2019.03.010.

Kunitomo H, Sato H, Iwata R, Satoh Y, Ohno H, Yamada K, Iino Y. 2013. Concentration memory-dependent synaptic plasticity of a taste circuit regulates salt concentration chemotaxis in Caenorhabditis elegans. Nature Communications 2013 4:1. 4(1):1–11. doi:10.1038/ncomms3210. [accessed 2024 Sep 17]. https://www.nature.com/articles/ncomms3210.

Lakowski B, Hekimi S. 1998. The genetics of caloric restriction in Caenorhabditis elegans. Proc Natl Acad Sci U S A. 95(22):13091–13096. doi:10.1073/PNAS.95.22.13091. [accessed 2024 Sep 17]. https://pubmed.ncbi.nlm.nih.gov/9789046/.

Lamitina ST, Morrison R, Moeckel GW, Strange K. 2004. Adaptation of the nematode Caenorhabditis elegans to extreme osmotic stress. Am J Physiol Cell Physiol. 286(4 55–4). doi:10.1152/AJPCELL.00381.2003/ASSET/IMAGES/LARGE/ZH00040418840005.JPEG. [accessed 2023 Apr 5]. https://journals.physiology.org/doi/10.1152/ajpcell.00381.2003.

Lamitina T, Huang CG, Strange K. 2006. Genome-wide RNAi screening identifies protein damage as a regulator of osmoprotective gene expression. Proc Natl Acad Sci U S A. 103(32):12173. doi:10.1073/PNAS.0602987103. [accessed 2023 Apr 5]. /pmc/articles/PMC1567714/.

Li L, Liu Z, Hu H, Cai R, Bi J, Wang Q, Zhou X, Luo H, Zhang C, Wan R. 2024. Dendrobium Nobile Alcohol Extract Extends the Lifespan of Caenorhabditis elegans via hsf-1 and daf-16. Molecules. 29(4):908. doi:10.3390/MOLECULES29040908/S1. [accessed 2024 Sep 17]. https://www.mdpi.com/1420-3049/29/4/908/htm.

Lin YF, Schulz AM, Pellegrino MW, Lu Y, Shaham S, Haynes CM. 2016. Maintenance and propagation of a deleterious mitochondrial genome by the mitochondrial UPR. Nature. 533(7603):416. doi:10.1038/NATURE17989. [accessed 2024 Nov 8]. https://pmc.ncbi.nlm.nih.gov/articles/PMC4873342/.

McKay JP, Raizen DM, Gottschalk A, Schafer WR, Avery L. 2004. eat-2 and eat-18 Are Required for Nicotinic Neurotransmission in the Caenorhabditis elegans Pharynx. Genetics. 166(1):161–169. doi:10.1534/GENETICS.166.1.161. [accessed 2024 Sep 17]. 10.1534/genetics.166.1.161.

Mendoza AD, Woodruff TK, Wignall SM, O’Halloran T V. 2017. Zinc availability during germline development impacts embryo viability in Caenorhabditis elegans. Comp Biochem Physiol C Toxicol Pharmacol. 191:194–202. doi:10.1016/J.CBPC.2016.09.007. [accessed 2024 Aug 30]. https://pubmed.ncbi.nlm.nih.gov/27664515/.

Moehle EA, Shen K, Dillin A. 2019. Mitochondrial proteostasis in the context of cellular and organismal health and aging. Journal of Biological Chemistry. 294(14):5396–5407. doi:10.1074/jbc.TM117.000893.

Mörck C, Olsen L, Kurth C, Persson A, Storm NJ, Svensson E, Jansson JO, Hellqvist M, Enejder A, Faergeman NJ, et al. 2009. Statins inhibit protein lipidation and induce the unfolded protein response in the non-sterol producing nematode Caenorhabditis elegans. Proc Natl Acad Sci U S A. 106(43):18285– 18290. doi:10.1073/PNAS.0907117106/SUPPL_FILE/0907117106SI.PDF. [accessed 2024 Sep 17]. https://www.pnas.org/doi/abs/10.1073/pnas.0907117106.

Mosley MC, Kinser HE, Martin OMF, Stroustrup N, Schedl T, Kornfeld K, Pincus Z. 2024. Similarities and differences in the gene expression signatures of physiological age versus future lifespan. Aging Cell. 00:e14428. doi:10.1111/ACEL.14428. [accessed 2024 Dec 17]. https://onlinelibrary.wiley.com/doi/full/10.1111/acel.14428.

Newell Stamper BL, Cypser JR, Kechris K, Kitzenberg DA, Tedesco PM, Johnson TE. 2018. Movement decline across lifespan of Caenorhabditis elegans mutants in the insulin/insulin-like signaling pathway. Aging Cell. 17(1). doi:10.1111/ACEL.12704. [accessed 2024 Aug 30]. /pmc/articles/PMC5770877/.

Ogg S, Ruvkun G. 1998. The C. elegans PTEN homolog, DAF-18, acts in the insulin receptor-like metabolic signaling pathway. Mol Cell. 2(6):887–893. doi:10.1016/S1097-2765(00)80303-2. [accessed 2024 Sep 16]. https://pubmed.ncbi.nlm.nih.gov/9885576/.

Paradis S, Ruvkun G. 1998. Caenorhabditis elegans Akt/PKB transduces insulin receptor-like signals from AGE-1 PI3 kinase to the DAF-16 transcription factor. Genes Dev. 12(16):2488–2498. doi:10.1101/GAD.12.16.2488. [accessed 2024 Sep 16]. https://pubmed.ncbi.nlm.nih.gov/9716402/.

Pincus Z, Mazer TC, Slack FJ. 2016. Autofluorescence as a measure of senescence in C. elegans: look to red, not blue or green. Aging. 8(5):889–898. doi:10.18632/AGING.100936. [accessed 2024 Sep 17]. https://pubmed.ncbi.nlm.nih.gov/27070172/.

Pohl F, Germann AL, Mao J, Hou S, Bakare B, Lin PKT, Yates K, Nonet ML, Akk G, Kornfeld K, et al. 2023. UNC-49 is a redox-sensitive GABAA receptor that regulates the mitochondrial unfolded protein response cell nonautonomously. Sci Adv. 9(44). doi:10.1126/SCIADV.ADH2584. [accessed 2024 Aug 30]. /pmc/articles/PMC10619936/.

Runkel ED, Liu S, Baumeister R, Schulze E. 2013. Surveillance-Activated Defenses Block the ROS–Induced Mitochondrial Unfolded Protein Response. Mango SE, editor. PLoS Genet. 9(3):e1003346. doi:10.1371/journal.pgen.1003346. [accessed 2021 Mar 31]. https://dx.plos.org/10.1371/journal.pgen.1003346.

Schaar CE, Dues DJ, Spielbauer KK, Machiela E, Cooper JF, Senchuk M, Hekimi S, Van Raamsdonk JM. 2015. Mitochondrial and Cytoplasmic ROS Have Opposing Effects on Lifespan. PLoS Genet. 11(2):1–24. doi:10.1371/JOURNAL.PGEN.1004972. [accessed 2024 Sep 17]. /pmc/articles/PMC4335496/.

Schulenburg H, Félix MA. 2017. The Natural Biotic Environment of Caenorhabditis elegans. Genetics. 206(1):55. doi:10.1534/GENETICS.116.195511. [accessed 2024 Sep 19]. /pmc/articles/PMC5419493/.

Shao L-W, Niu R, Liu Y. 2016. Neuropeptide signals cell non-autonomous mitochondrial unfolded protein response. Cell Res. 26(11):1182–1196. doi:10.1038/cr.2016.118. [accessed 2019 Jul 3]. http://www.ncbi.nlm.nih.gov/pubmed/27767096.

Sheng Y, Abreu IA, Cabelli DE, Maroney MJ, Miller AF, Teixeira M, Valentine JS. 2014. Superoxide dismutases and superoxide reductases. Chem Rev. 114(7):3854–3918. doi:10.1021/CR4005296/ASSET/IMAGES/MEDIUM/CR-2013-005296_0037.GIF. [accessed 2024 Sep 17]. https://pubs.acs.org/doi/full/10.1021/cr4005296.

Stefanello ST, Gubert P, Puntel B, Mizdal CR, de Campos MMA, Salman SM, Dornelles L, Avila DS, Aschner M, Soares FAA. 2015. Protective effects of novel organic selenium compounds against oxidative stress in the nematode Caenorhabditis elegans. Toxicol Rep. 2:961–967. doi:10.1016/J.TOXREP.2015.06.010.

Sulston JE, Schierenberg E, White JG, Thomson JN. 1983. The embryonic cell lineage of the nematode Caenorhabditis elegans. Dev Biol. 100(1):64–119. doi:10.1016/0012-1606(83)90201-4. [accessed 2024 Sep 19]. https://pubmed.ncbi.nlm.nih.gov/6684600/.

Traets JJH, van der Burght SN, Rademakers S, Jansen G, van Zon JS. 2021. Mechanism of life-long maintenance of neuron identity despite molecular fluctuations. Elife. 10. doi:10.7554/ELIFE.66955.

Urso SJ, Comly M, Hanover JA, Lamitina T. 2020. The O-GlcNAc transferase OGT is a conserved and essential regulator of the cellular and organismal response to hypertonic stress. PLoS Genet. 16(10):e1008821. doi:10.1371/JOURNAL.PGEN.1008821. [accessed 2022 Oct 31]. https://journals.plos.org/plosgenetics/article?id=10.1371/journal.pgen.1008821.

Urso SJ, Lamitina T. 2021. The C. elegans hypertonic stress response: big insights from shrinking worms. Cell Physiol Biochem. 55(Suppl 1):89. doi:10.33594/000000332. [accessed 2024 Aug 9]. /pmc/articles/PMC8265315/.

Wu Z, Senchuk MM, Dues DJ, Johnson BK, Cooper JF, Lew L, Machiela E, Schaar CE, DeJonge H, Blackwell TK, et al. 2018. Mitochondrial unfolded protein response transcription factor ATFS-1 promotes longevity in a long-lived mitochondrial mutant through activation of stress response pathways. BMC Biol. 16(1):147. doi:10.1186/s12915-018-0615-3. [accessed 2019 Dec 12]. http://www.ncbi.nlm.nih.gov/pubmed/30563508.

Yang W, Hekimi S. 2010a. Two modes of mitochondrial dysfunction lead independently to lifespan extension in Caenorhabditis elegans. Aging Cell. 9(3):433–447. doi:10.1111/j.1474-9726.2010.00571.x. [accessed 2020 Aug 12]. https://onlinelibrary.wiley.com/doi/full/10.1111/j.1474-9726.2010.00571.x.

Yang W, Hekimi S. 2010b. A Mitochondrial Superoxide Signal Triggers Increased Longevity in Caenorhabditis elegans. Tissenbaum HA, editor. PLoS Biol. 8(12):e1000556. doi:10.1371/journal.pbio.1000556. [accessed 2020 Aug 12]. https://dx.plos.org/10.1371/journal.pbio.1000556.

Yoneda T, Benedetti C, Urano F, Clark SG, Harding HP, Ron D. 2004. Compartment-specific perturbation of protein handling activates genes encoding mitochondrial chaperones. J Cell Sci. 117(18):4055–4066. doi:10.1242/JCS.01275. [accessed 2024 Sep 17]. 10.1242/jcs.01275.

Yu J, Gao X, Zhang L, Shi H, Yan Y, Han Y, Wu C, Liu Y, Fang M, Huang C, et al. 2024. Magnolol extends lifespan and improves age-related neurodegeneration in Caenorhabditis elegans via increase of stress resistance. Scientific Reports 2024 14:1. 14(1):1–17. doi:10.1038/s41598-024-53374-9. [accessed 2024 Sep 17]. https://www.nature.com/articles/s41598-024-53374-9.

Yu L, Yan X, Ye C, Zhao H, Chen X, Hu F, Li H. 2015. Bacterial Respiration and Growth Rates Affect the Feeding Preferences, Brood Size and Lifespan of Caenorhabditis elegans. PLoS One. 10(7):e0134401. doi:10.1371/JOURNAL.PONE.0134401. [accessed 2024 Dec 5]. https://journals.plos.org/plosone/article?id=10.1371/journal.pone.0134401.

